# Corollary discharge prevents signal distortion and enhances sensing during locomotion

**DOI:** 10.1101/2021.02.15.431323

**Authors:** Dimitri A. Skandalis, Elias T. Lunsford, James C. Liao

## Abstract

Sensory feedback during movement entails sensing a mix of externally- and self-generated stimuli (respectively, exafference and reafference). In many peripheral sensory systems, a parallel copy of the motor command, a corollary discharge, is thought to eliminate sensory feedback during behaviors. However, reafference has important roles in motor control, because it provides real-time feedback on the animal’s motions through the environment. In this case, the corollary discharge must be calibrated to enable feedback while avoiding negative consequences like sensor fatigue. The undulatory motions of fishes’ bodies generate induced flows that are sensed by the lateral line sensory organ, and prior work has shown these reafferent signals contribute to the regulation of swimming kinematics. Corollary discharge to the lateral line reduces the gain for reafference, but cannot eliminate it altogether. We develop a data-driven model integrating swimming biomechanics, hair cell physiology, and corollary discharge to understand how sensory modulation is calibrated during locomotion in larval zebrafish. In the absence of corollary discharge, lateral line afferent units exhibit the highly heterogeneous habituation rates characteristic of hair cell systems, typified by decaying sensitivity and phase distortions with respect to an input stimulus. Activation of the corollary discharge prevents habituation, reduces response heterogeneity, and regulates response phases in a narrow interval around the time of the peak stimulus. This suggests a synergistic interaction between the corollary discharge and the polarization of lateral line sensors, which sharpens sensitivity along their preferred axes. Our integrative model reveals a vital role of corollary discharge for ensuring precise feedback, including proprioception, during undulatory locomotion.

## Introduction

While moving through the environment, animals detect a mix of sensory signals originating from external sources and from the animal itself. Sensors provide feedback from the environment (exafference), but because they are embedded in a body moving through the environment, self-stimulation (reafference) is unavoidable. Reafference has commonly been considered a sensory artefact that interferes with appropriate behavior selection. For example, vigorous reafference could be mistaken for an impending attack, leading to the inappropriate activation of reflexive escape circuits (1, 2). Sustained reafference also fatigues sensors, reducing sensitivity and distorting the timing of neural signals (3, 4); in the extreme, overstimulated sensory circuitry may lead to excitotoxic cell death (5). For these reasons, many sensory pathways are modulated by feedforward neural signals activated in parallel with motor circuitry, which decrease sensor gain or eliminate reafferent signals altogether. These pathways, called corollary discharge (CD), mitigate negative consequences of reafference (1, 6). Well-studied examples of CD include the activities of inhibitory interneurons in crickets, crayfish, and tadpoles, which silence sensory circuitry during motor bouts (1, 2, 7, 8). This silencing is costly if it causes the organism to miss salient events in the environment. However, by focusing on sensing exafference and eliminating reafference, the field has tended to overlook the critical role of reafference in adaptive behavior and the negative consequences of suppressing reafference. Although reafference may entail negative consequences in terms of sensory ambiguity and sensor fatigue, it is nonetheless the error signal on which motor learning and sensor planning are based (1, 9–11). Reafference encodes crucial information about the animal’s own body parts with respect to the environment, such as relative orientation, speed, and acceleration (12–14). A key question of sensorimotor integration is therefore how CD can contribute to the regulation of the negative and positive aspects of reafference in light of behavioral demands.

The fish lateral line system is essential for informing behaviors such as schooling, prey detection, and rheotaxis (12, 15–18). The lateral line transduces fluid motions within the boundary layer through the bending of neuromast cupulae, depolarization of hair cells, and ultimately the transmission to the brain of afferent fiber spike activity (19–23). Because of this arrangement, the lateral line system is naturally subject to large amounts of reafference during undulatory swimming motions (23–25) (Figure 1 A). Fish have been found to rely on reafference signaling to provide proprioceptive feedback for swimming kinematics, though its mechanisms remain poorly understood [reviewed in (23, 24)]. The dependence on reafference becomes most evident at higher swim speeds and in turbulent flow conditions, in which cases the loss of lateral line feedback leads to progressively diverging kinematics (24, 26). It remains unclear what lateral line signals could provide proprioceptive feedback. Here, we reasoned that because the mechanical body wave (Figure 1 A) is consistently dysregulated by lateral line ablation (24, 26), sensing of the body wave itself is a component of proprioception. To understand how feedback from the lateral line thus contributes to kinematic control, we propose a model integrating the biomechanics of axial undulation (Figure 1 A) with lateral line neuromast biophysics. Axial undulations are fundamental to fish locomotion and are robustly transduced through periodic spiking in the lateral line (23–25, 27). The position of the peak of the body wave can therefore be inferred from the phase lag between the maximum spike rates of afferent fibers innervating sequential neuromasts (maximum cupular deflection, Figure 1 Aii,iii). As we show here, modulation of hair cell gain by the CD is a key component of the lateral line circuit that allows the fish to maintain a faithful representation of the body wave position.

**Figure 1.**
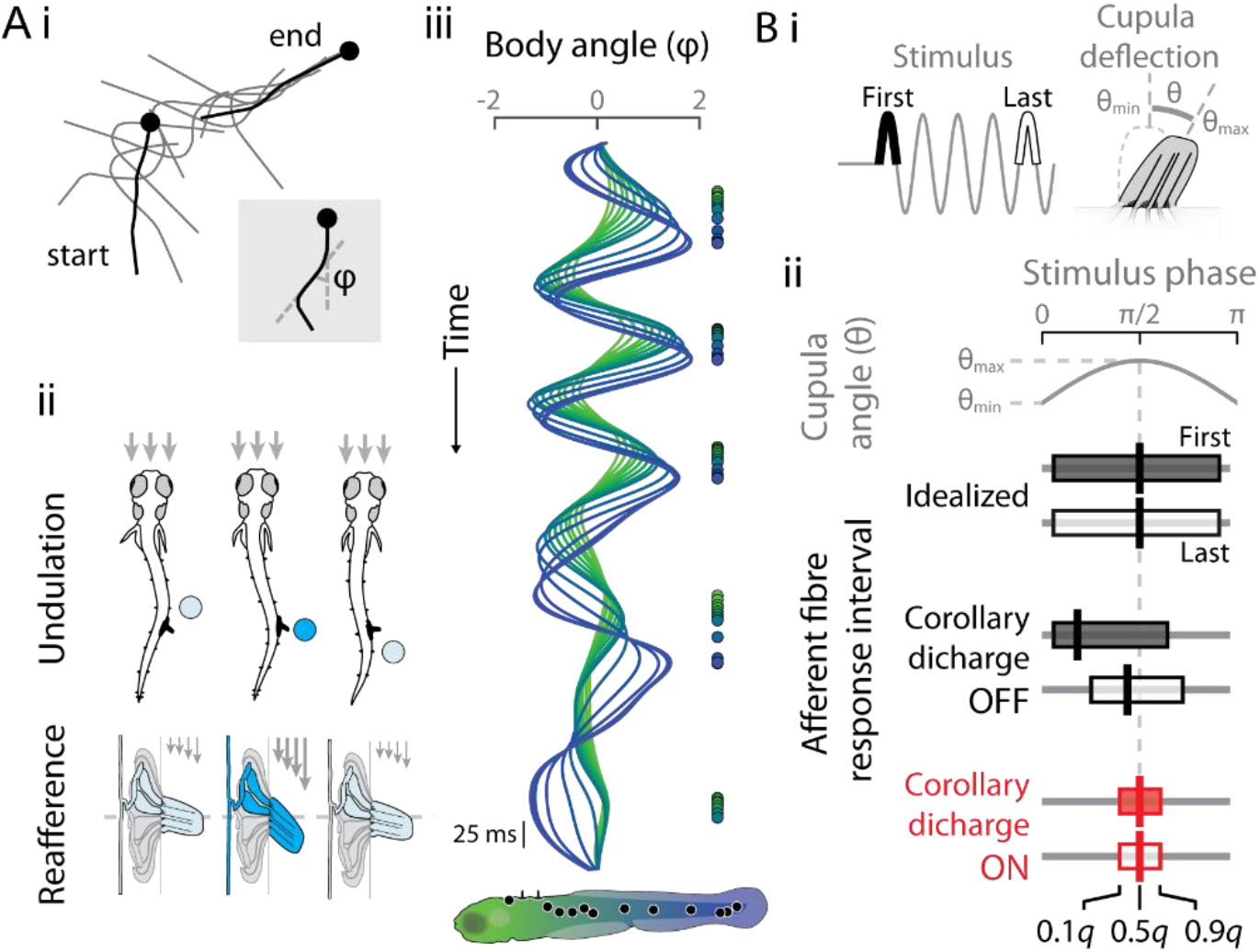
Timing of lateral line reafference during undulatory locomotion. (A) *i* Larval zebrafish swimming is characterized by body undulations. Body midline angles with respect to the head (ϕ) were determined by a spline from head (ball) to tail. *ii* Undulation involves a mechanical body wave that self-stimulates lateral line neuromasts embedded in the skin. Maximum cupular deflection and reafference (circles colored by reafferent intensity) are predicted to occur as the body wave peak passes the neuromast. *iii* Encoding peak timing allows the lateral line to signal the progression of the body wave by the phase lag between neuromast positions (green-blue color coding along body axis). (B) *i* Positional proprioception depends on faithful signalling of the maximum cupular deflection timing during repeated stimulation. *ii* An idealized sensor exhibits invariant responses that transduce deflection in the interval θ_min_ < θ ≤ θ_max_, such that peak responses coincide with the peak deflection phase, π/2. When the corollary discharge is OFF, habituation is predicted to cause phase-advanced responses with temporal distortions from the first to last stimulus presentation. Conversely, when corollary discharge is ON we predict that habituation is suppressed, and response intervals will be narrower, constant, and centered on π/2. In this way, gain modulation by the corollary discharge, together with the phase lag inherent to undulation, may enable positional proprioception through reafferent signalling in the lateral line system of fishes.

Hair cell physiology could be expected to limit the accuracy of sensory feedback during continuous stimulation. Hair cell systems, including the lateral line, exhibit high stimulus gain (nanometer accuracy, (28)) combined with finite vesicle populations (29, 30). Protracted stimulation, as occurs during locomotion, results in rapidly depressing responses, known as synaptic depression or habituation (21, 31–33), which can result in distortions of signal response phase, including delayed or advanced spike timings (3, 32). We illustrate this in Figure 1 Bii by comparing the spike phase distribution of an idealized sensor to that of a habituating sensor. The output of the idealized sensor is strictly proportional to cupular deflection along its axis of polarized sensitivity (34). The proportionality dictates that peak sensory feedback coincides with the maximum cupula deflection, which is taken to be at a stimulus phase of π/2 for our sinusoidal kinematic approximation. The median afferent fiber spike phase therefore serves as a detector of the peak of the body wave, and the phase lag between sequential neuromasts would mark its antero-caudal progression (Figure 1 A iii, B). Under the parameters of this model, a distortion of spike phase responses will impact the ability to accurately sense the body wave and coordinate kinematics. Synaptic depression leads to phase-advanced responses within a stimulus cycle (relative to the stimulus phase) and phase-delayed responses from the first to the last stimulus cycles [Figure 1 B, predictions based on (21, 32, 33)]. Further convoluting these distortions is the fact that lateral line habituation rates are highly variable, differing even among hair cells within a single neuromast (21). Heterogeneous habituation rates are expected to cause divergent afferent fiber signals over the course of the swim bout, complicating the task of tracking the timing of the body wave. Controlling heterogeneous gain within the lateral line should be of paramount importance in accurately sensing the position of the traveling body wave while swimming.

The CD to the lateral line tonically modulates the sensitivity of the lateral line during swim bouts (25, 35–37). Efferent projections from the hindbrain synapse on hair cells and evoke an inhibitory hyperpolarizing current while swimming, reducing the amount of presynaptic glutamate vesicle release (38–40). Hyperpolarization increases the current required to evoke exocytosis, which demands a larger cupular deflection to achieve the same afferent fiber activity. In addition to the reduced number of spontaneous and evoked afferent fiber spikes (25, 37, 40), the higher deflection threshold will also narrow the response interval (which in turn increases vector strength, Figure 1 Bii). The narrower interval results from delayed early spikes and advanced late spikes (as measured by the quantiles of the spike phase distribution), and affects sustained responses in two important ways. First, the propensity for habituation is reduced as the response interval narrows, because fewer vesicles are expected to be released (i.e., afferent activity is reduced). Second, preventing habituation will maintain afferent fiber spikes that are well-timed with respect to self-stimulation, and therefore can have proprioceptive functions. As the deflection threshold increases, it will eventually be equal only to the largest depolarization experienced by the hair cells in one neuromast, meaning that all afferent fiber activity will be concentrated on a single time point. We propose that this point is the maximum deflection of the cupula, which corresponds to the passing of the peak of the body wave (Figure 1 Aii). In this way, the self-stimulated afferent fiber activity signals peak body bending.

Because the lateral line sensors are so heterogeneous, how they are affected by CD may also be highly heterogeneous (41). This heterogeneity prompted us to study the role of CD on lateral line dynamics with three goals in mind. First, we use electrophysiology to detail among-unit differences in the magnitudes of habituation and CD on evoked spike rates. Second, we study afferent fiber responses, including spike rate and phase, when the CD is OFF (fish is inactive) or ON (fish is swimming). Finally, we simulate afferent fiber responses during CD OFF and ON to examine the lateral line’s capacity to accurately signal the timing of peak stimulation by the passing body wave. Our results are consistent with the hypothesis that CD to the fish lateral line evolved to precisely control hair cell gain while swimming and pass spikes that convey useful proprioceptive information.

## Results

### Interactions between habituation and corollary discharge affect spike rate and timing

The role of corollary discharge (CD) in regulating lateral line responses while swimming was examined in larval zebrafish (*Danio rerio*). Swimming activity was identified in immobilized larvae using extracellular recordings from ventral motor nerve roots, while lateral line activity was determined from patch clamp recordings (loose-patch configuration) of single afferent neurons (hereafter referred to as units) in the posterior lateral line afferent ganglion. Single neuromasts were sinusoidally stimulated using a glass bead affixed to a piezoelectric transducer at 5, 20, or 40 Hz to simulate the normal tail beat frequencies of freely swimming larvae (37, 42). Afferent fiber responses rapidly habituated to stimulation (exemplar trace at 20 Hz and peristimulus time histogram (PSTH), Figure 1 Ai, ii; 5 and 40 Hz in Supplementary Fig. S1; see also (21, 33)). The rapid decay followed by a refractory period (Figure 2 Aii) is similar to recordings in other recordings from lateral line and auditory afferent fibers (30, 33, 43). Responses to the first stimulus ranged from 1.2 – 5.4× that of the last stimulus. On average, the change in spontaneous and evoked spike rates (ΔS and ΔE, respectively) during swim periods (when the CD is active, CD ON) ~50% of inactive period (CD OFF; *p*<0.001, Supplementary Fig. S2; 5859 total swims), comparable to previous measurements (37, 40). However, the joint distribution of ΔS and ΔE revealed significant differences in some cells of the effects of CD on spontaneous and evoked spike rates (Figure 2 B *i*). We found that this was due to an interaction between the level of observed suppression and level of habituation (habituation = first-last response; ΔS × habituation: *p*=0.01, CI: - 0.54– −0.08). We therefore applied cluster analysis based on ΔS, ΔE, and habituation level to identify characteristics of different groups of responses among lateral line afferent units (Figure 2 B *i*). Four identified groups corresponded to each permutation of low and high levels of inhibition strength and degree of habituation, as verified by post-hoc analysis (Figure 2 B *ii*,*iii*).

**Figure 2.**
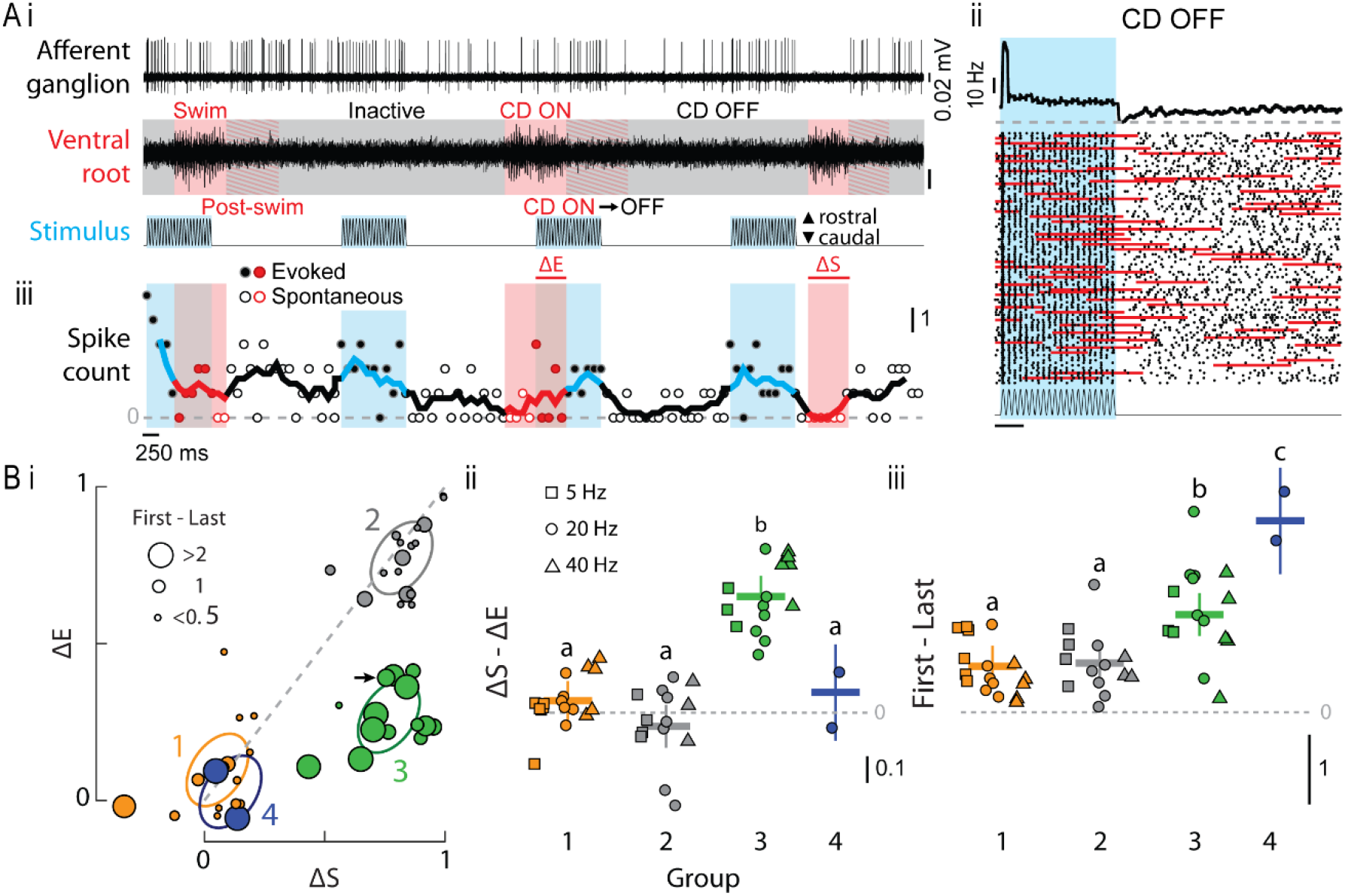
Afferent neuron spike activity in response to neuromast deflection during and between swims reveals interactions between habituation and corollary discharge (CD). (A) *i* Spontaneous and evoked lateral line afferent fiber activity during swim bouts (ventral motor root). Periods of swimming are highlighted in red, and periods of external stimulation of a single neuromast are highlighted in blue. Three periods were distinguished from ventral root recordings, corresponding to the activation of the CD: swim (CD ON), inactivity (CD OFF), and the post-swim transition between the two (CD ON⟶OFF). *ii* Peristimulus time histogram demonstrating spike rate (spike count per 100 ms) habituation when CD OFF, including decreased spike count and a post-stimulus refractory period. *iii* The impact of swim bouts (CD ON) on spontaneous (open face) and evoked (closed face) responses were analyzed within highlighted intervals. In this trace, the relative decrease in spontaneous spike rate (ΔS) was greater than that of the evoked spike rate (ΔE). There was an immediate restoration of evoked spike activity after the swim bout ended. (B) *i* Correlation of ΔS and ΔE reveals distinct groups of afferent responses at all frequencies (0=no suppression, 1=complete suppression) that depends on the level of habituation (First – Last spike count). Arrow highlights record corresponding to the trace in A. *ii*-*iii* Response groups were identified by cluster analysis of ΔS, ΔE, and level of habituation. Post-hoc comparisons among groups show inequality of ΔS and ΔE (*ii*) and degree of habituation (*iii*). 5 Hz: n=13 afferent neurons from N=12 individuals; 20 Hz: n = 22, N=21; 40 Hz: n=15, N=14.

The differences among groups suggested that interactions between habituation and CD identify a significant source of heterogeneity within and among afferent fibers during swim bouts. In order to understand how this heterogeneity could impact reafference signalling in our model (Figure 1), we next examined the dynamics of afferent spike count and phase in response to stimulation during CD OFF (black in Figure 3). The PSTHs reveal substantial phase distortions from the first to last stimulus intervals at each frequency (Figure 3 A). In parallel to the decline of spike count, the phase distortions resulted in narrower response intervals over time, and therefore increased vector strength (Figures 3 Bi, Ci; quantified through generalized additive models, GAM). The inverse relationship between spike count and vector strength was likewise observed among the four cell groups, such that Groups 1 and 2 exhibited lower average spike counts and vector strengths compared to Groups 3 and 4. To quantify time-dependent changes in lateral line sensitivity, we calculated afferent fiber gain (20) as the product of spike count and vector strength. Based on this measure, we suggest that cells in Groups 1 and 2 cells have low gain (average gain <1) whereas units in Groups 3 and 4 cells have high gain (gain >1; Figure 3 Ciii). Average and group-specific gain strongly declined due to habituation (Figure 3 Biii). The gain of units in Groups 1 and 2 was slightly greater than unity for initial stimuli, but fell to substantially less than unity subsequently. Units in Groups 3 and 4 maintained gain ≥1 but with very large relative changes (Figure 3 Biii). The PSTHs also suggested that habituation more substantially delayed the timing of the earliest spikes (spike latency) than of later spikes. We corrected for the frequency-dependent phase shift caused by linear conduction delays in the afferent fibers (20) by subtracting each unit’s median phase at each frequency. All quantiles of the spike phase distribution were delayed, indicating that the response interval is shifted as well as being narrowed (as indicated by vector strength).

**Figure 3.**
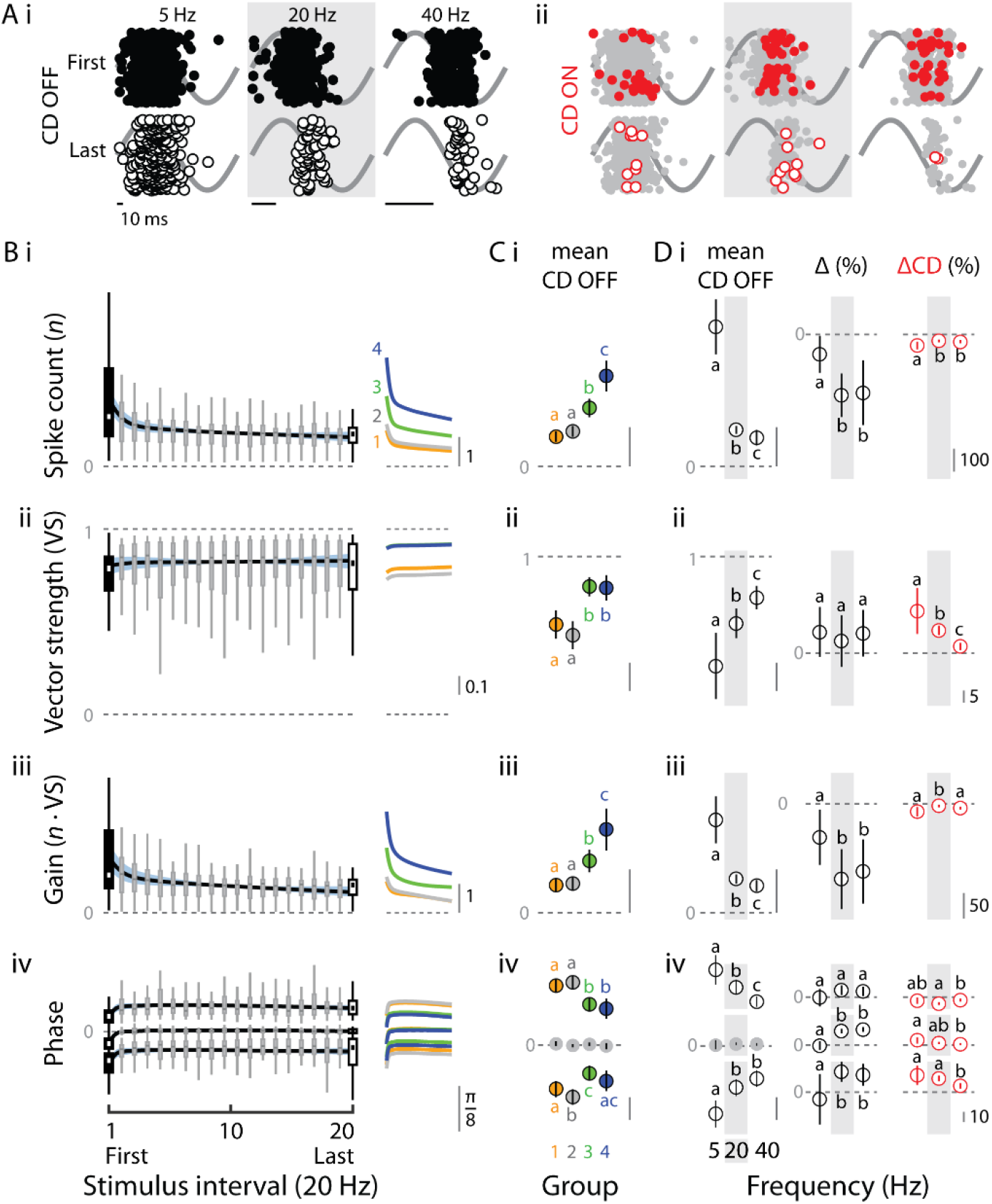
Effects of habituation and corollary discharge (CD) on evoked afferent fiber spike responses. (A) PSTH of spike phases (circles overlaid on sinusoidal stimulus) grouped by the first (solid symbols) and last (open face) stimulus period and by CD OFF (*i*) or CD ON (*ii*). *i* At all frequencies, CD OFF results in habituation and significantly delayed response intervals. *ii* CD ON reduces spike number and causes narrowed and shifted responses intervals with respect to CD OFF (shaded for comparison). Data for individual in Figure 2 A. (B) Box-and-whisker plots of stimulus-averaged spike count, vector strength, gain, and response phase at 20 Hz. The average time-dependent changes between first and last intervals during CD OFF were quantified by smooth functions. *Right*: Time courses of responses for response groups identified in Figure 2. (C) Mean responses among groups. Group 1 and 2 cell responses were not statistically distinguishable, and included lower mean evoked spike rates, vector strength, and gain compared to Group 3 and 4 cells. Note that median phase cannot be analyzed, as described in the text. (D) Frequency-dependence of mean responses, habituation, and CD ON. Higher stimulation frequencies resulted in reduced spike counts, higher vector strength, and lower gain. Narrower response intervals coincide with the change in vector strength. The relative effects of habituation. The relative impact of habituation (Δ%) on spike count and gain was lower at 5 Hz compared to 20 and 40 Hz, whereas the relative effect of CD (ΔCD%) was relatively greater. Across frequencies, there was an equal but nonsignificant overall increase in vector strength with habituation, whereas the relative effect of CD decreased with higher frequencies. There was no observable effect of habituation to the 5 Hz stimulus on spike phase, but CD caused a relatively large delay of the earliest spikes and relatively small advance of the late spikes, resulting in later median phases. These effects diminished at higher frequencies, reducing the change in median phase. Sample sizes as in Figure 1. *p*<0.001 for depicted smooth functions in all panels. All effects in C and D are means and standard errors.

Average responses among groups coincided with the differences in vector strength, such that cells in Groups 3 and 4 had much narrower response distributions (Figure 3C iv). Average responses varied widely across frequencies. Spike count and vector strength were inversely related, respectively decreasing and increasing at higher frequencies, resulting in a decline in per-stimulus gain to less than unity (Figure 3 Di-iii). The narrower response intervals are reflected in the response phases (Figure 3 Div). Because average responses varied widely, we expressed the effect of habituation as the difference between the first and last stimulus intervals relative to the average CD OFF response (Δ% in Figure 3). The effect of habituation was not explicitly parametrized in our models, so was calculated from samples drawn from the model posterior distributions (the smooth functions are nonparametric). Habituation had a small effect on spike count and gain at 5 Hz compared to 20 and 40 Hz (Figures 2 Bii, Cii). There was no statistically distinguishable effect on vector strength across frequencies, which also all overlapped zero. However, response phase was significantly delayed at 20 and 40 Hz, but not at 5 Hz (Figure 2 Div). Moreover, the phase of early spikes was more affected than that of late spikes; for example, at 20 Hz, 0.1*q* and 0.5*q* phase were delayed by 0.48π (CI: 0.23–0.73) and 0.44π (CI: 0.36–0.51), whereas 0.9*q* was 0.28 (CI: 0.04–0.39). This asymmetry implies a narrower response interval which, as noted, was only weakly supported by the vector strength measurement. This could be due to differences in calculation and statistical power between the two measurements; vector strength is calculated from all spikes, including outliers that fall outside the preferred phase (which can be seen in e.g., Figure 2A). The use of quantiles eliminates these outliers, which we suggest helps to identify consistent changes despite the highly variable effects of habituation.

We next sought to quantify the average effects of CD ON. The relative difference between CD OFF and CD ON was modelled as an intercept shift between the average responses in each condition, expressed relative to CD OFF (ΔCD% in Figure 3). Across all frequencies, CD ON decreased spike count and increased vector strength, including at 5 Hz despite the negligible effects of habituation. This is superficially similar to habituation, but in contrast to habituation, CD delayed early spike phases but advanced late spikes’ phase (Figure 3 Div). In this case, the asymmetric change in response intervals implied by the spike phase quantiles was supported by the vector strength, particularly at 5 Hz. We interpret these results to indicate that both habituation and CD OFF decrease sensors’ gain for the stimulus, which progressively reduces lateral line sensitivity to all but the cupular peak deflection (stimulus phase of π/2). Thus the effect of intense habituation, like CD ON, should be to create a more faithful representation of the body wave, as implied by our model in Figure 1. However, the change in gain due to habituation will depend on highly variable intercellular differences (Figure 2 B,C) as well as variable effects of motor frequency and duration (see also, (37, 42)). Thus, habituation should be an intrinsically unreliable mechanism for increasing the fidelity of afferent responses with respect to the input stimulus. It is also noteworthy that the responses to habituation and CD were more similar at 20 and 40 Hz than 5 Hz. We suggest that this is because 5 Hz stimulation lies below typical larval tail beat frequencies (37, 42), and therefore levels of habituation, to which the CD is adapted.

### Tuning hair cell gain prevents habituation and maximizes post-swim sensitivity

Our recordings suggested that CD can prevent habituation to maintain sensitivity to post-swim stimuli. We typically observed an immediate restoration of sustained evoked spike rates when a swim bout ended during a stimulus interval (Figure 2 A), or even bursts of evoked spikes otherwise only seen at the start of the stimulus presentation (examples highlighted in Supplementary Fig. S1). Swim-timing dependent responses in spike rate and phase are obscured by our open-loop experimental setup and subsequent analyses (Figure 2) in which we averaged over evoked responses occurring at different relative times within distinct swim bouts. Thus, we normalised all swim bouts and an equal post swim period to unit duration and then analysed response properties (spike rate, vector strength, gain, and spike phase quantiles) of swims falling entirely within or outside the stimulus period. This allowed us to quantitatively address the role of CD during and after swim bouts, and our prediction that CD preserves constant phase response.

Afferent unit responses during and after swim bouts substantially differed from responses over the course of habituation. In contrast to the decline in spike count due to habituation, evoked spike counts increased nearly linearly over the swim duration (significance of smooth, *p*<0.001, Figure 4 A). Whereas spike activity was depressed after the stimulus period with CD OFF (Figure 2 A *ii*), spike activity after the swim bout actually peaked. The time course of spontaneous spike rate likewise exhibited a post-swim peak (*p*<0.001; Figure 4 A), which is consistent with the effect of electrical stimulation of inhibitory efferent fibers that causes vesicle accumulation and then post-inhibition rebound activity (38). The increase in spontaneous spike rate and evoked spike count over time can therefore be traced to the increasing cumulative probability of release over time due to vesicle accumulation (38). That it occurs during stimulation points to the CD protecting the vesicle pool from evoked release, given the level of stimulus that we applied.

**Figure 4.**
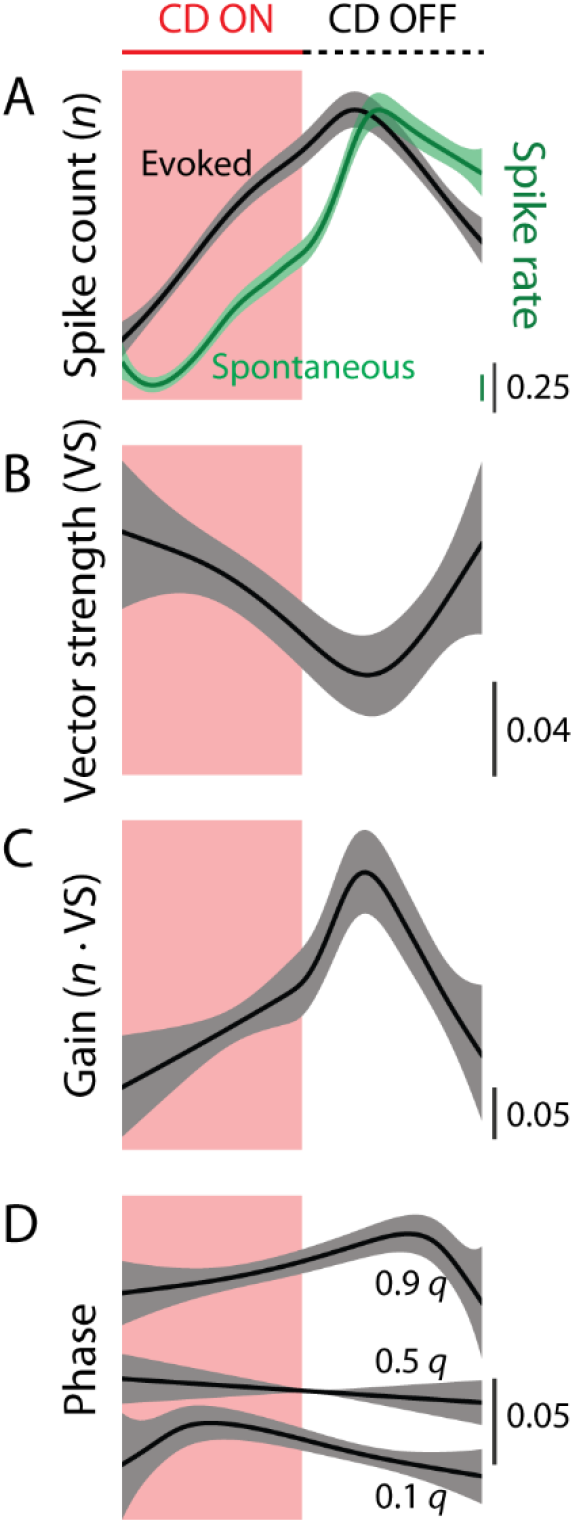
Corollary discharge counteracts habituation and preserves sensitivity after swimming. Swim bouts and a subsequent post-swim period of equal time were normalized to unit duration. These periods correspond to CD ON (swimming, red) followed by CD OFF (white), although residual effects of CD ON persist into the post-swim period (37). All effects normalised to start of trace. (A) Spontaneous (green) and evoked (black) spike counts were maximal in the post-swim period. The rise in evoked spike count with CD ON contrasts to the fall in responses with CD OFF (see Figures 2 Aii, 3 B). Significance of smooth functions, *p*<0.001. (B) Vector strength was initially high, representing a narrow response interval, and exhibited a post-swim minimum (*p*<0.001). (C) Afferent signal gain reached a maximum during the post-swim period (*p*<0.001). There appears to be an inflection point toward increased gain corresponding to the end of CD ON. (D) The response interval width is narrower during CD ON, as as indicated by a smaller separation between 0.1*q* and 0.9*q*. During CD ON, 0.1*q* is significantly delayed (rising trend) but becomes advanced (falling trend) through the swim and CD OFF periods (0.1*q*: *p*=0.02; 0.9*q*: *p*=0.04). In contrast, 0.9q becomes progressively delayed to a peak in the CD OFF period, but then substantially advances. This suggests that response intervals are narrow during CD ON but widen with CD OFF. A relatively constant median response phase (0.5*q*, *p*=0.33) was preserved throughout the CD ON and OFF intervals.

We predicted that CD ON would maintain a constant response interval and median spike phase from the first to last stimulus periods (Figure 1 Bii). Instead, vector strength somewhat decline over the course of the swim (*p*=0.005), reaching a post-swim minimum coinciding with the timing of the evoked spike count maximum (Figure 4 B). The regulation of spike rate and interval width by CD resulted in maximum gain (spike count × vector strength) for stimuli in the post-swim period (Figure 4 C), indicating a preservation to respond to post-swim stimuli. The changes in response interval over time were due to continuous changes in the timing of both the earliest and latest spikes. With CD ON, the response interval was narrow due to delayed 0.1*q* and advanced 0.9*q*, relative to the post-swim phase with CD OFF (0.1*q*: *p*=0.02: 0.9*q*, *p*=0.008). Although the phase quantile dynamics were asymmetrical, an essentially constant median spike phase (*p*=0.31, Figure 4 D) was preserved throughout.

### Corollary discharge prevents phase distortion due to habituation

Our results indicated that variation in gain is fundamental to understanding heterogeneity within the lateral line, and thus the impact of CD on sensing. We developed a computational simulation of afferent fiber responses to further probe the interactions between heterogeneity in afferent fiber gain, CD strength, and afferent fiber response phase. Our simulation is based on a conceptual model of vesicle pool depletion (CD OFF) or conservation (CD ON). Simulated vesicles transitioned between four states: bound and ready for release; releasing, which was taken as a spike; released and unavailable for signaling; or regenerating by returning to the ready state (Figure 5B *i*). Simulated fibers recapitulated the time-averaged features of the CD OFF condition, namely a strong response to the first stimulus, a habituated response to subsequent stimuli, a refractory period after stimulation, and variable response phase during stimulation (Figure 5 A ii, *iii*). We applied this model to examine how the strength of inhibition affects the behavior of simulated afferent fibers with rapid first responses and strong habituation (high gain and low regeneration rates), or slow first responses with no habituation (low gain and high regeneration rates; Figure 5 B *i*). Following from Figure 2, inhibition strength was equal to a 0, 50, or 90% decrease of simulated spontaneous spike rate (ΔS). We then examined the steady evoked spike rate (ΔE) relative to the habituated rate (depicted in Figure 5 A*ii*).

**Figure 5.**
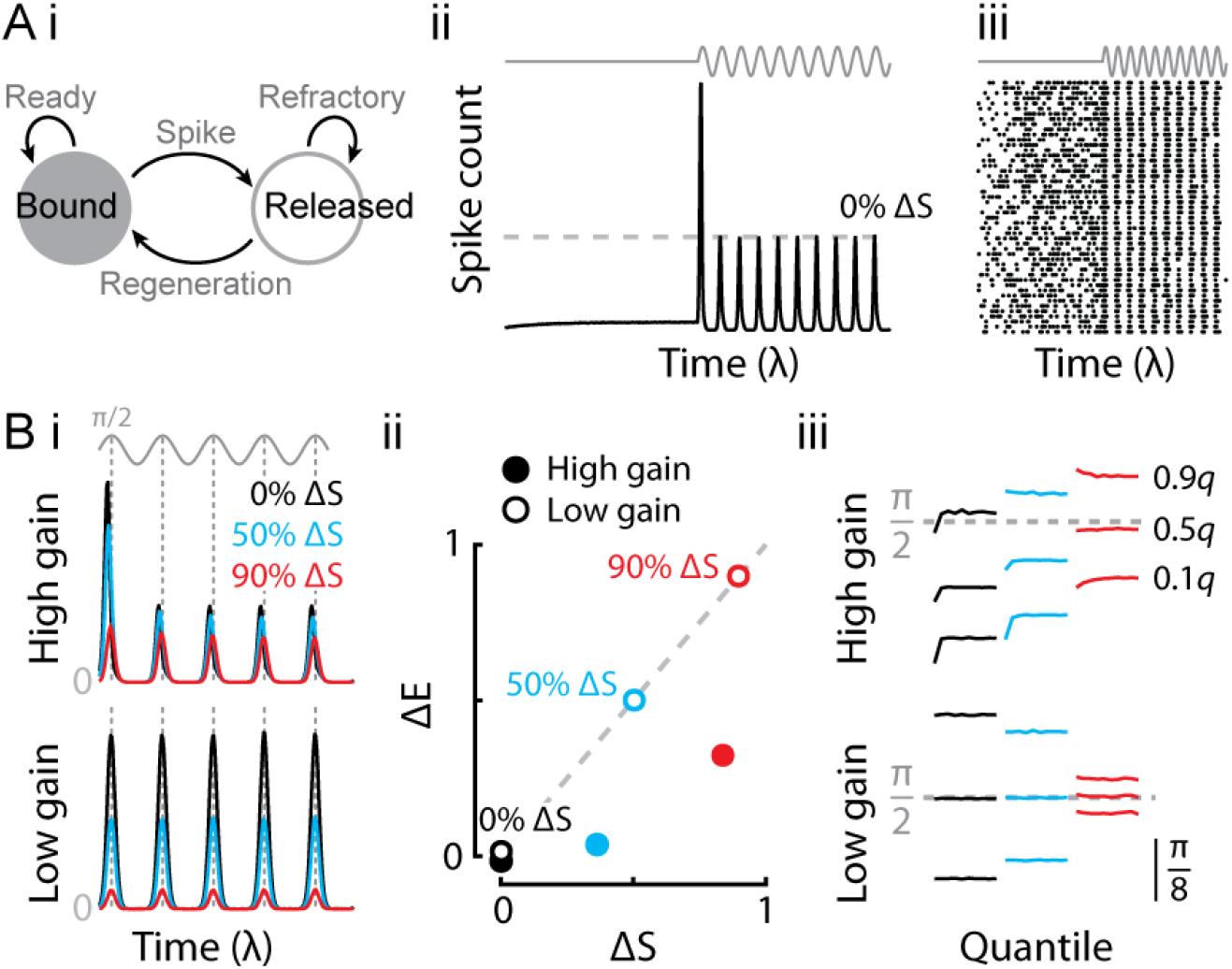
Corollary discharge counteract habituation to increase proprioceptive information content of afferent signals. (A) *i* The relationship of afferent fiber spike phase to the peak of reafference was examined through a state transition model reflecting simple vesicle dynamics. Simulated vesicles may be bound and ready for release, or they may transition to a released state (a spike) and become unavailable until regenerated (transition back to the bound state). *ii* Average responses of a simulated fiber without suppression (0% ΔS) demonstrate major features of experimental recordings including strong initial and subsequently weaker responses, and a distinct refractory period after stimulation. The average habituated level is taken as the reference level for further comparisons (dashed line). *iii* PSTH of responses in *ii* showing variable spike timing within individual stimulus presentations. (B) *i* Differential effects of corollary discharge on simulated afferent fiber responses. Vertical dashed lines denote the stimulus peak timing at π/2. We simulated the effects of inhibition equal to 0, 50, or 90% ΔS, on an afferent fiber with strong habituation (high gain) or without (low gain). *ii* Correlation of simulated ΔS and ΔE reveals divergent effects of suppression on high- and low-gain parameter sets. Points at the origin are staggered for clarity. *iii* The high-gain kinetics exhibited time-dependent phase-leading behavior, whereas the low gain kinetics had constant response intervals centered on the stimulus peak at π/2. When the high-gain kinetics were strongly inhibited (90%), the median response phase converged on the stimulus peak at π/2, without time-dependent distortions.

The effects of simulated CD on spike rate differed markedly depending on the level of habituation (Figure 5 B *i*). Inhibition had much less effect on the high-gain (habituating) compared to low-gain (non-habituating) parameter set. The relationship between ΔS and ΔE mirrored that observed experimentally (Figure 2 B) in that the level of habituation predicted a divergence in the apparent level of inhibition by CD. We found ΔE≪ΔS in the high-gain parameter set and ΔE=ΔS in the low-gain set (Figure 5 B *ii*). The responses were identical at low levels of suppression, regardless of habituation rate, which supported the identification of Group 4 highly-habituating but lowly-suppressed cells (Figure 2 B).

Finally, we examined how CD affects phase relationships of simulated responses with respect to the peak stimulus. Simulated high gain caused responses that were phase-leading with respect to the stimulus input (peak at π/2), but became delayed over the course of habituation (Figure 5 B *iii*). Conversely, the low-gain responses exhibited constant phase centered on the peak of the input stimulus, in fact similar to our idealized sensor (Figure 1 Bii). In both parameter sets, moderate inhibition (50%) had minimal effects on spike rate or phase, which might explain why the strength of suppression appears to be bimodally distributed (40). However, strong inhibition (90%) suppressed the tendency of the high-gain parameter set to habituate, as shown by the relative spike rates of Figure 5 Bi). In addition, strong inhibition suppressed the distortion of the median phase by spreading spikes more evenly over the first interval and maintaining constant phase thereafter. The same level of inhibition greatly narrowed response intervals of the non-habituating parameter set. Crucially, high levels of inhibition in both parameter sets resulted in median phase approaching the stimulus peak. We conclude that CD functions to regulate the temporal structure of afferent spikes to achieve constant median phase with zero or minimal delay.

In our experimental recordings, the majority of strongly habituating units (Group 3 in Figure 2 Bi) were strongly inhibited, as judged by ΔS. Our simulations support labelling the Group3 units as high gain. Strong inhibition by CD will prevent habituation by moderately reducing the signalling interval, and cause their median response phase to converge on π/2. In contrast, strong inhibition of weakly-habituating units (Group 2, Figure 2 Bi) dramatically reduces their response interval, which remains centered on π/2. Altogether, we expect that the CD should maintain consistent feedback that can signal the position of the mechanical body wave (Figure 1).

## Discussion

Axial undulation is the ancestral mode of vertebrate locomotion, predating paired appendages, and is defined by a traveling body wave that enables propulsion through complex environments. To stabilize this mode while navigating, undulating fish must have a sense of self-motion and body position, yet they lack many of the classes and much of the distribution of proprioceptors found in other vertebrates (44, 45). Fish nonetheless possess a complement of proprioceptors which appear to be suited to sensing undulation-associated stresses from the inside out: within the body, on the surface of the body, and at the body-fluid interface. This integrated sensing system is supported by mechanosensors within the spinal cord (46), stretch sensors in the skin (45), and fluid-coupled sensing in the lateral line (24). The role of the lateral line has been the least clear, in large part because an external proprioceptor would fundamentally differ from better-understood mechanisms of actuator- and joint-embedded proprioceptors, like stretch receptors (44). Moreover, the lateral line is not a dedicated proprioceptor, and so it has been uncertain how apparent motion artefacts could be discriminated from environmental stimuli and then transformed to useful proprioceptive feedback. We develop a model based on sensing the alternating flow currents within the undulating body’s boundary layer (Figure 1), including a vital role for corollary dischage (CD). We hypothesize that fish robustly detect the peak of neuromast cupular deflection (Figure 1), but that intrinsic sensor heterogeneity will increase response variability to obscure the timing of peak deflection (Figure 1, 3). We find that CD reduces variability while swimming by suppressing habituation (Figure 4) and, through stronger suppression of high-gain cells identified in this paper, homogenizes response phases (Figure 5). By suppressing habituation, the CD maintains robust sensing after the swim bout (Figure 3), when a fish could otherwise experience a period of sensory blindness.

Differences in gain can serve as a major source of heterogeneity in lateral line initial and sustained sensitivity (Figures 2, 3). Maintaining populations of cells with large variation in response gain ensures a wide dynamic range for functions like frequency and spatial filtering (21, 22, 47). For instance, high-gain sensors would provide an advantage for early sensing of a predator’s flow perturbations (48). The wide range in gain may provide sensory advantages for a stationary animal, but would also increase variability in response to self-stimulation and thus complicate the problems of reafference while swimming (Figures 3, 5). Units with greater gain exhibit earlier vesicle release during the stimulus cycle and experience greater habituation, which depletes vesicle pools until they are regenerated. This in turn decreases sensitivity and results in temporal distortions (Figures 3, 5). This is particularly evident in units from response Groups 1, 2, and 3, which have initial gain greater than unity but fall to ≤1 thereafter (Figure 3 Bi). The experimental and computational results identify differences in average phase and changes in phase due to habituation according to gain (Figures 3, 5).

The variability in afferent responses caused by self-stimulation during swimming leads us to posit that differential gain modulation through CD is a crucial element of sensorimotor integration. The interaction between level of habituation and CD strength gave rise to four identifiable groups of cells (Figure 2) that were subsequently identified in our simulations as low- and high-gain units with weak or strong suppression (Figure 5). Across all cell groups, hair cell hyperpolarization by the CD prevented habituation (Figures 2, 5) and sustained sensitivity throughout the swim bout and after (Figure 4). Simulated respones of low- and high-gain afferent units revealed that strong CD was required to prevent the habituation of the high-gain units. This may explain why nearly all the highly habituating units also had strong inhibition (i.e., those in Group 3, Figure 2). A small subset of cells, labelled Group 4, were high gain but with weak inhibition (Figure 2 B, 5 Bii). We were unable to determine each unit’s directional preference, but the weak inhibition might suggest rostrocaudal polarization (40). If so, then their high gain notwithstanding, these units would be expected to experience only weak reafference and habituation during forward swimming. In addition to controlling habituation among units with variable gain, the CD also largely homogenizes responses with respect to the stimulus phase, ensuring consistent timing of feedback throughout the swim bout (Figures 4 C, 5 Biii). Importantly, the CD ensures that the response interval is narrow and centered on the peak stimulus phase (Figures 1 B, 5 B *iii*). Interestingly, this implies that the CD synergistically interacts with hair cell polarization to sharpen directional tuning, a possibility that had been dismissed on the grounds that the efferent system is not sufficiently specific (25). However, our model shows that the synergy emerges from a hyperpolarization mechanism rather than any degree of efferent specificity.

Given that strong suppression was necessary to control habituation in the high-gain units, why might all hair cells not be suppressed as strongly as possible? Our findings support the hypothesis that CD evolved to suppress afferent spiking as much as necessary, but not more (40). One consequence of universally strong CD inhibition would be the elimination of weaker reafference signals from retrograde flows that are relatively weaker during forward swims (40). A second consequence is that strong inhibition would lead to post-swim artefacts. In the absence of stimulation, CD ON caused post-swim rebound spiking (Figure 4), similarly to galvanic stimulation of cholinergic fibers in the auditory system (38). Excessive inhibition causing rebound spiking in the caudal to rostral direction could be misinterpreted as a predator attack from behind. A requirement of the CD is therefore that it be calibrated to control motion artefacts without introducing new ones.

We propose that the CD is an essential component of proprioceptive feedback in most fishes. It is, though, conspicuously absent in sea lamprey, a basal jawless fish that have been shown to rely on lateral line proprioception for kinematic control (24). Biofluiddynamic differences between adult lamprey and larval zebrafish notwithstanding, we suggest that the sea lamprey’s proprioceptive lateral line can operate without CD because of its low self-stimulation frequency. Typical adult sea lamprey tail beat frequencies lie between ~2-4 Hz (24); at 5 Hz stimulation in zebrafish, we observed minimal habituation but the CD had substantial effects including delayed spike latency and median spike phase (Figure 3 D). At low frequencies, the lateral line therefore behaves similarly to our idealized sensor (Figure 1 Bii), which would suggest that suppression through CD may be deleterious to sensing. In this respect, it is important to consider that hair cells are hyperpolarized for half the stimulus cycle when they are deflected against their preferred direction. This is critical because it not only enables in directional specificity, as is most frequently studied, but it also provides a vesicle regeneration interval *within* the swim cycle and not just after. Indeed, this may be the major reason why we observed little habituation at 5 Hz stimulation, despite the very large number of spikes per stimulus interval (Figure 3A, D). A consequent prediction is that there should be minimal suppression in slow or sedentary species that nonetheless possess a CD, because they will experience minimal habituation. Indeed, the oyster toadfish (*Opsanus tau*) is a teleost that possesses a CD but exhibits very little suppression in spike rates during slow forward swimming (27). This echoes the dynamics that we observed and simulated in the zebrafish lateral line, which is that strong inhibition is principally active in cells with greater propensities for habituation. Altogether, this suggests that the CD is crucial for fast swimmers with higher tail beat frequencies, and that there is a wide evolutionary scope for its strength and role in feedback. Fishes are enormously diverse, so the strength and role of CD in feedback should be studied within their varied ecological and biomechanical contexts.

We propose that controlling the effects of habituation is crucial, but we also recognize that some degree of habituation that does not impair sensing can be advantageous. Habituation can enable functions like source localization through bilateral comparisons or other detectors, as occurs in the auditory and electrosensory systems (3,4, 30). In a related manner, habituation could contribute to rheotactic behaviors, in which fish orient upstream by comparing bilateral differences in water velocity, presumably encoded by differences in spike rate (12). Bilateral differences in flow will cause differential habituation, and as a logical extension of our results, bilateral differences in spike timing as well. Temporal information, as well as rate, could therefore be involved in orienting to flows. Overall, the processing of temporal information must occur on real behavioral time scales much shorter than the tens of seconds or even minutes studied in previous lateral line work (20, 49, 50). Exploratory swim bouts of zebrafish larvae last as few as 3-5 tail beats [150-250 ms at 20 Hz, (42)], so reafference and habituation dynamics over a few tail beat cycles represent a substantial portion of the swim. Rheotactic swims last longer, up to 15 tail beats [750 ms at 20 Hz, (51)], so dynamics on a longer time scale are also relevant to sensing capacity during and after sustained behaviors. Our stimulus period of 1 s represents the upper limit of fictive swim durations (Supplementary Fig. S2). An essential advance will therefore be to understand the course of hydrodynamic stimulation and CD’s inhibition on realistic behavioral time scales, because the strengths of stimulation and inhibition change both during and after swims (25, 35, 37, 41, 42). Time-dependent changes in inhibition strength (41) may control more intense reafferent signals at the swim start that are associated with stronger first tailbeats (42), rostral-caudal flow (40), or the kinetics of the rapid vesicle pool (52). Precisely and faithfully preserving the phase characteristics of a stimulus is a key function of auditory (30). We argue that through the actions of the CD, the lateral line likewise faithfully signals the phase lag between sequential neuromasts to provide proprioceptive feedback of the mechanical body wave during undulatory swimming.

## Materials and Methods

### Electrophysiological recordings

Adult zebrafish (*Danio rerio*) were obtained from a laboratory-bred population and housed as described previously (e.g., (20, 37, 53), in accordance with protocols approved by the University of Florida’s Institutional Animal Care and Use Committee. Protocols were based on our previous study (37). Larvae from 4 to 6 dpf were paralyzed by immersion in 1 mg/mL α-bungarotoxin (Reptile World Serpentarium, St Cloud, Florida) diluted in 10% Hank’s extracellular solution (37), and then washed and replaced in extracellular solution until experiments (>30 mins). α-bungarotoxin blockade of the neuronal-type nicotinic acetylcholine receptor (nAChR) α9 found in neuromasts is reversible after a brief washout period (54). Larvae were pinned laterally to a Sylgard-bottomed dish using etched tungsten pins inserted through the dorsal notochord at the anus, at a second location immediately caudal to the yolk sac (approximately at neuromast L1), and through the otic capsule. The preparation was then covered in extracellular solution for recording. Fish health was monitored by blood flow.

We reliably obtained single-unit afferent recordings, or occasionally two clearly differentiable units, using borosilicate glass pipettes with diameters ~15 μm. At this pipette diameter, it was necessary to approach the ganglion under high pressure (–150 to –200mmHg) in order to break in. Once a unit was located, pressure was released sufficiently to maintain a stable recording (typically between 0 and –75 mmHg). In most cases, the afferent recording was stable for hours. Afferent recordings were performed at 20 kHz with 50V/mV gain, Bessel filter of 1 kHz, and an AC high-pass filter of 300 Hz. Signals were amplified through an Axon Instruments CV-7B head stage (MD: Molecular Devices, San Jose, CA) and an Axoclamp 700B amplifier (MD), then recorded with a Digidata 1440A digitizer and pClamp10.7 (MD). Spike times were obtained by thresholding in Clampfit 10.7 (MD). Once an afferent unit was identified, we systematically probed neuromasts for the stereotypical afferent evoked response to a periodic stimulus (20), using a glass bead attached to a single-axis piezo stimulator (30V300, piezosystem jena, Inc., Hopedale, MA). Dipole fluid motions of this type are commonly used to assess near- and far-field responses to oscillatory stimuli, such as from prey or conspecifics (12, 48, 55). The dipole vibrating at 20 or 40 Hz simulates neuromast stimulation by the larva’s own movements, because reafference *in vivo* is associated with alternating (23, 24) rather than direct currents (40, 41). Stimulation at 5 Hz was also performed to compare responses to a frequency at the extreme lower limit of typical tail beats (42). The probe head was roughly spherical with a diameter of 56.6 μm, formed by melting the tip of a recording pipette using a MF-830 microforge (Narishige International, Amityville, NY). Probing started at D1 and continued down the body until the evoked response was elicited. The majority of units could be stimulated by D1 (n=16) or D2 (n=3), and were classified as D-type, and only a few units did not exhibit responses until we reached L1 (n=1), LII.1 or LII.2 (each n=2), or LII.3 (n=1), classified as L-type. In one case we were unable to locate any attached neuromast. Note that, because we only identify the most proximally innervated neuromast, the afferents could have been connected to more distal neuromasts as well (22). After identifying the neuromast, the probe was placed one diameter away from the cupula, and stimulation was performed with a constant ~8 μm amplitude. At no point was the cupula itself touched, which substantially alters responses of the hair cell-afferent group (20).

Once an afferent and associated neuromast were identified, we obtained extracellular recordings of the ventral root (VR) arborizations in the myotomal cleft (37). Pressure in the pipette was manually controlled through a syringe. Ventral root recordings were acquired as per afferent units, but with a gain of 100V/mv. Swim bouts were preprocessed by thresholding the VR signal in Clampfit 10.7, and then manually curated using custom software in R 3.6.0 (56). Ventral root recordings were stable for ~15–60 mins. All swims were spontaneous or elicited by a light flash.

### Data processing

We recorded 100 sweeps with the stimulus on for 1 s then off for 2 s, which ensured recovery of afferent spike rate in all but one possible case (Figure 3 B). We first examined responses at 20 Hz (*n*=22), then one or both of 40 Hz (*n*=15) or 5 Hz (*n*=13) depending on preparation stability. Additional recordings were performed as required to guarantee sufficient numbers of swims for analysis. Traces were divided into intervals according to the presence or absence of a swim or stimulation period (Figure 2). The post-swim interval was equal in duration to the swim (37). Spontaneous spike rates were calculated from periods in which none of these conditions were present. Evoked responses were quantified either as a function of stimulus number or the average over the stimulation period (Figure 2). Within a stimulus interval of period *T*, the phase *θ* of a spike occurring at time *t* was expressed as *θ*=*t*/*T*. The phase can then be used to examine distributional properties of spikes, such as central tendency or width. The commonly reported vector strength (VS, e.g., (20)) for a set of *n* spikes is calculated by,

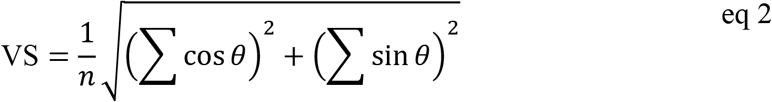

and is related to the circular variance according to *σ*^2^ = −2log (VS) but is constrained to lie within [0,1], with VS→1 as the interval narrows. Single spikes’ information content is quantified by the gain, which is calculated by the product of spike count and VS. We additionally computed lower, median, and upper quantiles (0.1, 0.5, and 0.9 quantiles, *q*) of the phases, which reflect the tails of the spike phase distribution while being more robust than the minima (i.e., latency to first spike) and maxima.

Spike counts were modelled with a quasi-Poisson distribution, and arcsin-transformed vector strength and the spike phase quantiles were modelled through Gaussian distributions. The intermittent swims resulted in fewer intervals with swims than inactivity, and so less precise estimates of responses during the swim period. For example, an evoked spike might be observed in 85 of *N*=95 (*p*=0.89) stimuli presentations while inactive, but four of *N*=5 (*p*=0.80) presentations while swimming, an apparent decrease in probability during the swim. Uncertainty due to sampling was incorporated through observation weights of 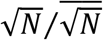. In total, we observed 5859 swims across all individuals and frequencies. Modelled relationships were transformed to the response scale for plotting.

### Statistical analysis

Differences due to habituation and swimming were characterized through generalized linear models (GLM). Modelling was performed with the R function *glmer()*, with Tukey post-hoc tests performed with the function *glht()* in package *multcomp* (57). These contrasts were chosen to examine average differences in each condition and to compare our results to literature values (Supplementary Fig. S2). Significant differences in means were assessed by confidence intervals that excluded 0. All models were fit by maximum likelihood. Throughout, we provide relevant estimate effects and 95% confidence intervals; full statistical models (R summaries), including relevant *p*-values, are included in SI Appendix. Statistical significance was tested at α=0.05.

The relationships between ΔS, ΔE, and habituation were explored through linear models and cluster analysis. The level of habituation was estimated as the difference in responses to the first and last stimulus. We found that directly averaging first and last responses was prone to high variance, likely due to sampling error, and therefore we used a more stable estimate obtained from the predicted response at each stimulus for each individual at each frequency in the population GAM (Figure 3 C, see below). The prediction of ΔE from ΔS and habituation was then studied in a linear mixed effects model (R package *nlme*, (58)), with individual nested in frequency. The slope was forced through zero so that the approximate confidence intervals of the predictors could be compared to zero or unity to assess significance. Individual ΔS, ΔE, and habituation responses were clustered using finite Gaussian mixture models (59), and post-hoc analyses of the groups performed to illustrate the magnitude of differences between clusters.

We used generalized additive models (GAMs) to model the nonlinear responses over the stimulus period as a non-parametric smooth function (60, 61). We employ an adaptive spline basis set (61) with the upper limit of spline knots respectively set to the frequency, i.e., the number of response intervals. We allow for random variation in the function among identified cell response groups (factor-smooth, (61)), which allows for response heterogeneity among classes (e.g., (21, 22)). A fixed effect of intercept was included, as well as random slopes between stimulus intervals and individual neuron. The smooth portion of the fit is centered on zero, so differences in average responses between spikes labelled as occurring during inactivity or swimming are accounted for through parametric intercept differences (inactivity was the reference level). The relative difference in intercepts, expressed as swim intercept/inactivity intercept, is used to model the average change during swims. Arcsin-transformed vector strength was modelled with a Gaussian error distribution, and spike counts and gain were modelled with a quasi-Poisson distribution. Lower quantile (0.1*q*) was cube-transformed prior to analysis to stabilize the variance.

The GAM modelling is also appropriate for examining changes in evoked responses during and after the swim. Within each individual at each frequency, we selected swims that fell entirely within (1650 swims) or outside (3348) the stimulus interval, which were then aligned by their terminus and normalized to unit duration. We analyzed responses from the swim start up to one swim duration after the swim end as a function of relative time, using an adaptive spline basis. Differences among stimulus frequencies were included as a fixed effect and a factor smooth frequency allowing for frequency-specific trends (61). A random intercept term was included for individual cells. Spontaneous activity was averaged over time (spike rate) rather than per stimulus period.

We sought to compare responses between frequencies. The average differences between CD OFF and ON are parametrized in the model, but the smooth fit modelling habituation is nonparametric. We therefore compare model predicted responses by performing 1000 repeated draws from the posterior distributions of the statistical models of first and last stimulus periods, and of the average CD OFF and ON responses. From these draws, we calculate the predicted average effect of habituation (last-first response) and corollary discharge (CD ON-CD OFF). This process results in means and standard errors of each quantity of interest. We compare the probability that the resulting distributions differ at α=0.05.

### Time-inhomogeneous Markov chain simulations

We explored potential causal mechanisms of the phase relationships discovered experimentally by simulating different biophysical parameter sets giving rise to strong or weak afferent fiber habituation. Hypothetical vesicle release underlying afferent fiber spiking was modelled as a transition between state 1 (a bound vesicle) and state 2 (released vesicle, recorded as a spike). The discrete-time transition matrix is given by,

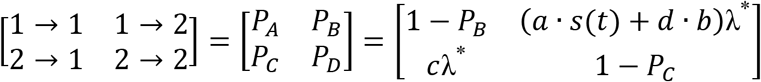

The transition matrix elements are normalized to stimulus wavelength divided by sampling frequency (λ*, simulated at 1000 Hz). Spontaneous spike rate, *b*, is the number of expected spikes per stimulus wavelength. The magnitude of inhibition is given by the expected suppression of the spontaneous spike rate, ΔS. In the absence of inhibition (*d*=1), a vesicle has non-zero release probability for all *a*=0. The stimulus is modelled by a time-dependent stimulus function, *s(t)*, multiplied by the gain, *a*. When *s(t)* is constant, a time-invariant stimulus, a rapid burst of evoked spikes rapidly settle into a constant rate that reflects the balance of regeneration and release probabilities. For time-varying stimuli, the responses depend on the relative coefficient magnitudes. The expected regeneration rate per λ* is given by *c*. Habituation can occur when regeneration rates are outpaced by the high probability of release (*a*≫*c)*. A parameter set for high-probability release vesicles was chosen to qualitatively reflect the readily releasable pool in hair cells and retinal bipolar neurons (29, 62) (*a*=0.04, *b*=4, *c*=4). Conversely, the low-probability parameter set was chosen to minimize habituation through low release rates and high regeneration (*a*=4, *b*=40, *c*=0.04). The lower *b* in the latter case was chosen to prevent spontaneous spike rates from dominating the evoked responses. The stimulus duration was 20 λ*, with *s*=0 for the first half of the period to allow vesicles to regenerate. Inhibition was applied only during the stimulus period, with *d*=0, 0.5, or 0.1 (0, 50, or 90% inhibition). Simulations were run for 2-20k iterations as required to depict smooth responses in Figure 5. The first two iterations were discarded as burn-in to avoid impacts of the initialization conditions. We then analyzed averaged responses from each parameter set.

Data and modelling code have been deposited in the online repository available at DOI 10.6084/m9.figshare.13034012.

## Acknowledgments

We are extremely grateful to Dr. Noah J. Cowan for numerous insightful discussions, and Drs. Yuriy Bobkov, Paula Duarte Guterman, Miriam H. Richards, James A. Strother, and anonymous referees whose critical evaluation greatly improved the manuscript. This research was supported by National Institutes of Health (DC010809), National Science Foundation (IOS1257150, IOS1856237), private foundation grants to JCL, and support from the Whitney Laboratory for Marine Biosciences to DAS.

## Classification

Biological Sciences, Neuroscience

## Author Contributions

DAS, ETL, and JCL conceived experiments; DAS and JCL designed experiments; DAS performed experiments and analyzed results; DAS wrote the manuscript; DAS, ETL, and JCL edited the manuscript. Resources: JCL; Project administration: JCL; Funding acquisition: JCL. All authors approved the manuscript in its final form.

