## Supplementary Figures for "Corollary discharge prevents signal distortion and enhances sensing during locomotion"

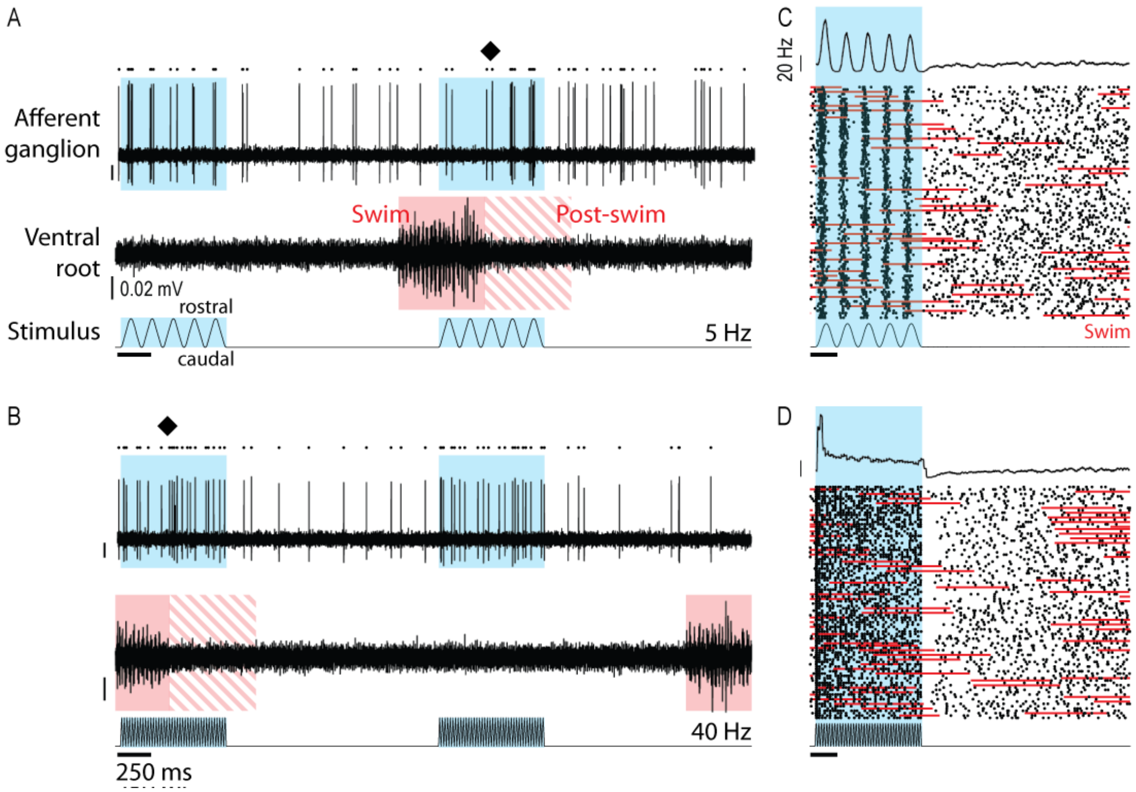


**Figure S1.** Exemplar traces of responses to 5 and 40 Hz stimulation, same individual as Figure 1 A. Evoked spike rates rebound after the end of swimming (diamonds).

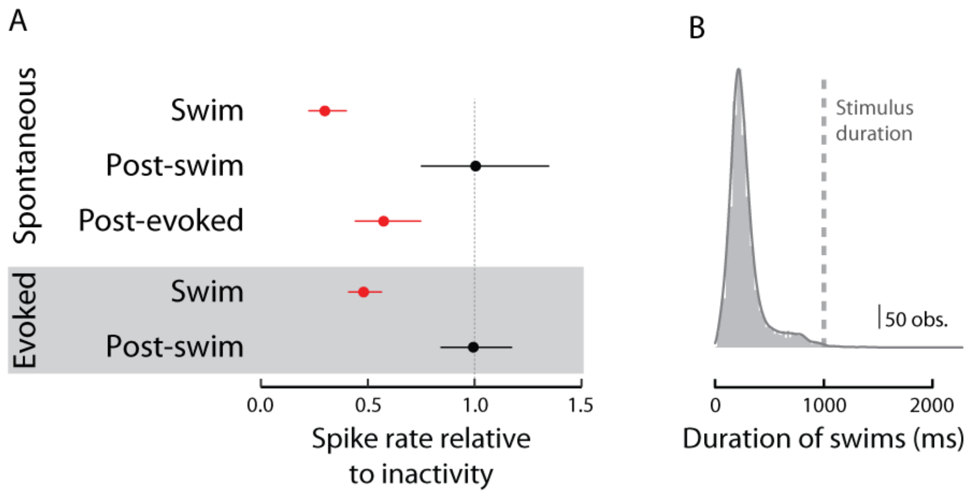


**Figure S2.** Average responses during swims. A) Mean and 95% confidence intervals of spontaneous and evoked spike rates in the swim, post-swim, and post-evoked periods, relative to spontaneous or evoked spike rates while the fish is inactive. $\Delta S$ and $\Delta E$ were calculated as one minus this relative level, and therefore $\Delta S = 1-0.3 = 0.7$ (confidence intervals, CI: 0.60-0.78) and $\Delta E=0.52$ (CI: 0.43-0.59). Changes in post-swim spike rates averaged over one swim duration (Figure 3) do not significantly differ from unity. Spike rates were significantly reduced after the stimulus period, relative to subsequent spontaneous spike rates (Figure 1 A). B) Distribution of fictive swim durations in this study. Most swims are <500 ms, corresponding to the distribution of real swims (1, 2). The stimulus duration of 1 s encompasses the upper limit of swim durations.
